# Aggregation is a context-dependent constraint on protein evolution

**DOI:** 10.1101/2021.05.10.443436

**Authors:** Michele Monti, Alexandros Armaos, Marco Fantini, Annalisa Pastore, Gian Gaetano Tartaglia

## Abstract

Solubility is a requirement for many cellular processes. Loss of solubility and aggregation can lead to the partial or complete abrogation of protein function. Thus, understanding the relationship between protein evolution and aggregation is an important goal. Here, we analysed two deep mutational scanning experiments to investigate the role of protein aggregation in molecular evolution. In one data set, mutants of a protein involved in RNA biogenesis and processing, human TAR DNA binding protein 43 (TDP-43), were expressed in *S. cerevisiae*. In the other data set, mutants of a bacterial enzyme that controls resistance to penicillins and cephalosporins, TEM-1 beta-lactamase, were expressed in *E. coli* under the selective pressure of an antibiotic treatment. We found that aggregation differentiates the effects of mutations in the two different cellular contexts. Specifically, aggregation was found to be associated with increased cell fitness in the case of TDP-43 mutations, as it protects the host from aberrant interactions. By contrast, in the case of TEM-1 beta-lactamase mutations, aggregation is linked to a decreased cell fitness due to inactivation of protein function. Our study shows that aggregation is an important context-dependent constraint of molecular evolution and opens up new avenues to investigate the role of aggregation in different cellular contexts.

## I. INTRODUCTION

The majority of proteins function as monodispersed ordered species dissolved in intra- or extra-cellular aqueous fluids. There are, however, abnormal or pathologic states in which proteins aggregate, often becoming insoluble and dropping out of solution [1, 2]. In other cases, protein assembly leads to a liquid-liquid phase separation producing a reversible soluble coacervate [3, 4]. A liquid-to-solid phase transition is often detrimental to the cell as it is associated with a loss of the function that the protein has evolved to attain in the specific context [5]. In some cases, aggregates could acquire aberrant functionalities, which can have consequences for the onset and development of disorders [6]. Although these principles are valid for a large part of the proteome [7, 8], aggregation can be required in specific cases to increase the functionality of a biological system. This occurs, for instance, for Pmel17, a protein found in mammalian melanosomes, whose aggregation is necessary for the production and storage of melanin [9]. Given the double valence of protein aggregation that can result either in the formation of toxic assemblies or physiologically required organelles, it has long been debated what could determine whether aggregation is beneficial or detrimental from an evolutionary perspective.

Here, we set to clarify this complex matter and question of whether aggregation is an important constraint in protein evolution that can be the discriminant between beneficial and detrimental situations by considering two cases: we studied molecular evolution of proteins in an endogenous versus exogenous host. One of the important factors that reduce the danger of aggregation is in fact the presence of interaction partners that protect the regions that would otherwise be involved in self-assembly [10, 11]. Indeed, the interfaces through which proteins interact are often highly aggregation prone, so that their engagement in interactions reduces propensity to self-assembly [12]. This means that regions under the evolutionary pressure of protein function may be the same that lead to aberrant aggregation [13–15]. Any situation, such as mutations or anomalous expression, could result in disruption of these beneficial interactions with consequent aggregation and depletion of normal function.

Expression of a protein in a host different from the natural one is of course an artificial case but represents a prime example in which natural interactions cannot occur. This situation could induce a completely unnatural environment in which proteostasis (or maintainance of protein homeostasis) is impeded [16]: if not promptly degraded, an exogenous protein could establish aberrant interactions and aggregate [3, 17].

To evaluate the effect of exogenous versus endogenous expression, we analysed two large-scale data sets in which variants of an exogenous and an endogenous protein were expressed in different host systems. We focused on molecular evolution data of human TAR DNA-binding protein 43 (TDP-43) exogenously expressed in yeast and bacterial TEM-1 beta-lactamase endogenously expressed in *E. coli*. TDP-43 is involved in various steps of RNA biogenesis and processing, including alternative splicing [18]. TEM-1 beta-lactamase is an enzyme that hydrolyzes the beta-lactam bond in susceptible beta-lactam antibiotics, conferring resistance to penicillins and cephalosporins in bacteria [19]. The mutational library of TDP-43 was designed to introduce mutations specifically in the intrinsically unstructured region near the C-terminus of the protein (290–331 or 332–373). The library of beta-lactamase mutants targeted instead the whole stably structublack enzyme.

With these two systems, we investigated whether and how aggregation acts as a constraint of the evolutionary process by finely tuning the selection of proteins to fit in a specific environment [8]. While aggregation leads always to protein depletion, we aimed to determine if the detrimental or beneficial effects sensitively depend on the context and nature of the selection sieve.

## II. MATERIAL AND METHODS

### A. Random Mutations and Algorithms

In our analyses, we used mutation of TDP-43 and TEM-1 beta-lactamase sequences that have the same length as their respective wild types. Each amino acid was selected for a mutation with the same frequency. Changing the amino acid frequencies from uniform to weighted according to the natural occurrences does not change the results (Supplementary Material). As for the negative sets used in TDP-43 and TEM-1 beta-lactamase analyses, the amount of random mutations was matched with the experimental occurrences of the positive sets.

Predictions were carried out with Zyggregator [2]. This algorithm estimates whether a protein aggregates into solid-like aggregates (fibrillar or protofibrillar structures) on the basis of combinations of physico-chemical properties such as hydrophobicity, helical and beta-propensities [20]. Since the physico-chemical properties of the amino acids change with the environment in which the aggregation processes take place, the coefficients in the Zyggregator algorithm were fitted on a database of experimental rates collected for homogeneous conditions and the predictions were only performed in the vicinity of such conditions. Here, we used a linearized version of Zyggregator [21] to speed up the analysis of the large-scale sets of mutants. By linearization we mean a procedure in which the aggregation propensity was computed considering the amino acid composition of aggregation-prone regions predicted by Zyggregator. This calculation transforms each amino acid and close neighbours (7 amino acids as defined in previous works [2, 21]) in a numerical value to estimate the aggregation propensity.

The camFOLD method [22] was used to calculate the folding propensity profiles of proteins according to a model that exploits the same physicochemical properties exploited by Zyggregator but combined in a different way.

### B. Data generation description

The experimental datasets were retrieved from two publications in which error-prone oligonucleotide synthesis was adopted to comprehensively generate the mutational data sets of human TDP-43 [23] and TEM-1 beta-lactamase [24].

Briefly, human TDP-43 was expressed in *S. cerevisiae* [23]. Deep sequencing before and after TDP-43 mutant library induction was used to quantify the relative effects caused by each variant on growth (the input culture was split into two cultures and fitness was calculated from changes of output to input variant frequencies relative to wild type). After quality control and filtering (three replicates of deep mutational scanning experiments, minimum mean input read count threshold of ten) [23], the approach quantified the toxicity of 1,266 single and 56,730 double amino acid changes with high reproducibility.

TEM-1 beta-lactamase was expressed in *E. coli* and twelve libraries of variants were generated [24]. These libraries underwent cell phenotypic selection in a growth functional assay to retain only the variants coding for a functional protein. Differently from TDP-43, only positive fitness variants were sequenced. The number of transformants were in the same order of magnitude of the sequencing capacity of the next-generation sequencing platform (between 100 thousand and 1 million) to guarantee good library complexity. The different generations (or evolutionary cycles) were produced serially, which indicates that the number of mutations increase with respect to the wild-type. The sequencing was performed at the 1st, 5th and 12th generation. While in the 1st generation the dose of antibiotics for phenotypic selection was at 25ug/mL, in the 5th and 12th generation the level was raised to 100 ug/mL to allow stricter selection. In the last generation library a total of 177,253 protein variants with the same length of the wild-type protein (i.e., 286 amino acids without inclusion of premature stop codons) were retrieved[24].

### C. Data Analysis

For the analysis of the TDP-43 data set, we used both the negative and positive fitness scores measured relatively to the toxicity of wild-type protein in *S. cerevisae*. By contrast, in the TEM-1 beta-lactamase case, we only had variants selected through the evolutionary process and comparison with an antagonistic set was not directly possible. To carry out the analysis, we generated a negative set that was as close as possible to the pool of mutations in the experiments of deep mutational scanning. We tested the randomized negative set using the TDP-43 case: we designed a set of sequences with one or two random mutations depending on the original composition of the pool and performed a comparative analysis. Our results indicated similar performances using a negative or random set, which suggests that they are interchangeable. Once the procedure to generate the random set was tested, we proceeded to build a negative set for the TEM-1 beta-lactamase case. We designed a number of mutations following the rule of keeping the number of mutations present in the positive set. To avoid statistical biases, we analyzed only the sequences from the positive set that have the same length of the wild-type. We analyzed the 1st, 5th and 12th generation sets for which sequencing data were available. To quantify how aggregation changes along the data sets, we computed the Receiver Operating Characteristics (ROC) curves and their relative Area Under the Curve (AUC) for each sub-data set using the corresponding random set as a negative control. We measured the AUC vs the number of mutations keeping track of the evolutionary cycles. Some sub-data sets with specific mutations were poorly populated in the later cycles (as the number of mutations increase in each evolutionary cycle) and therefore could not be used for our statistical analysis. To avoid the issue and maintain consistency, we analyzed only sub-data sets that had a size at least 5% that of the largest sub-data set at each evolutionary cycle. We then tested how aggregation and folding propensities changed while a protein accumulated random mutations. We generated a control data set selecting mutations randomly from the 20 natural aminoacids. At each step of evolution, we ran the linearised Zyggregator algorithm to get a score to quantify the aggregation and folding propensities. We did this for all the variants and checked if the sequences increased or decreased their propensities at each random mutation.

## III. RESULTS

### A. Aggregation and folding predictions

We first tested if a linearized version of the Zyggregator approach could be used to perform proteome-wide analyses, which comes with the advantage of having faster calculations (see Materials and Methods).

As shown in Figure 1, the linearized version of the algorithm shows a Pearson correlation of 0.78 with the original Zyggregator algorithm. This result indicates that the linearization procedure is able to capture the fundamental features of the Zyggregator algorithm and can be used in our analysis.

**FIG. 1:**
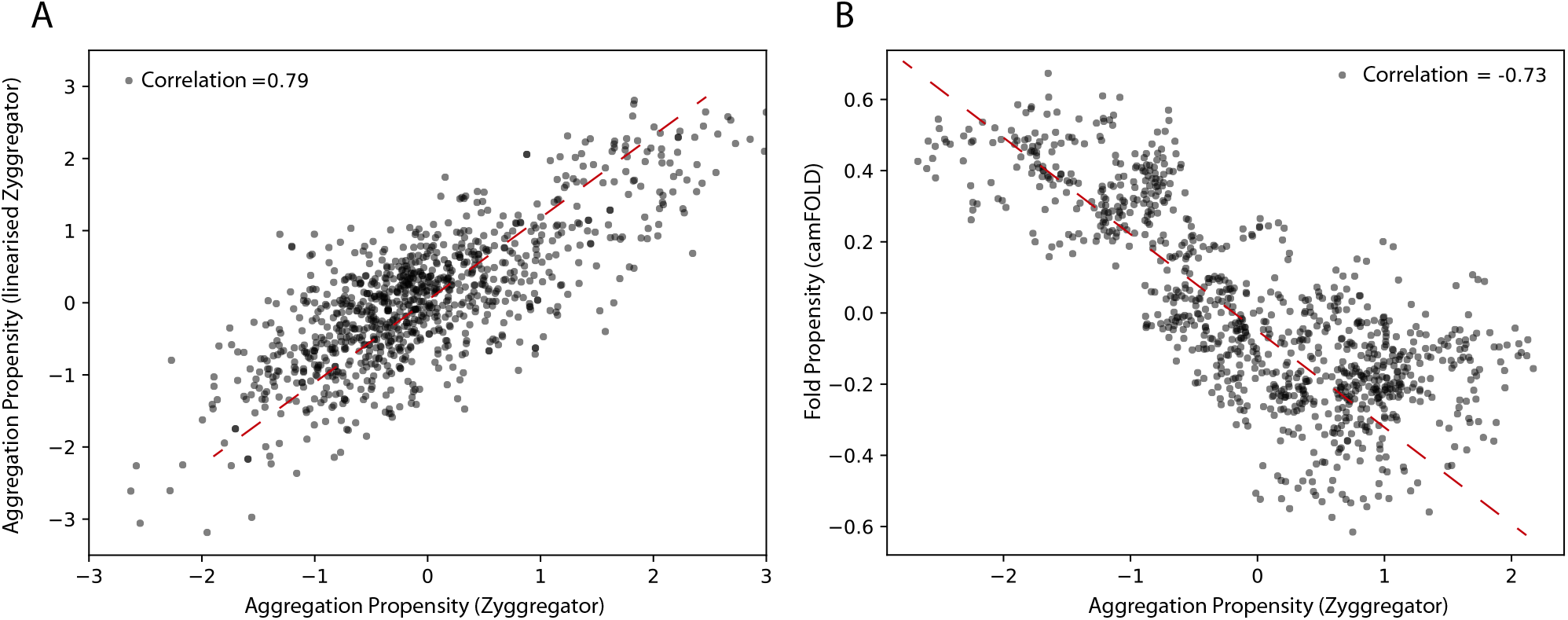
Aggregation and folding propensity of the human proteome. A) Scatter plot of Zyggregator algorithm and ‘linearized’ version computed on 5000 random proteins from the human proteome. The correlation among the two methods is 0.79, which justifies the use of the linear approximation of Zyggregator. B) Scatter plot of Zyggregator and camFOLD scores. Folding and aggregation propensities are anti-correlated with a Pearson correlation coefficient of -0.73.

In agreement with previous work [22], we found that the Zyggregator and camFOLD scores (see Material and Methods) are functionally related for a large set of proteins taken from the human interactome (Figure 1, panel B; correlation of -0.73). In other words, proteins with a high propensity to fold show poor aggregation propensity.

### B. Random mutations increase the aggregation propensity

Can evolution drive sequences to aggregate ? Clearly much depends on the context and functions of the proteins of interest, but it is possible to make some general considerations.

When we mutated sequences in the human proteome, the aggregation propensity rose and the folding propensities decreased (Figure 2, panels A and B). This result indicates that in many cases, including enzymes acting in aqueous solutions [25, 26], protein function requires a stable native fold. This is in agreement with the hypothesis that proteins “live on the edge” due to opposing effects on their sequences caused by the evolutionary pressure to prevent aggregation at the concentrations needed by the cell and the random mutational processes that inherently increase the aggregation propensities [8].

**FIG. 2:**
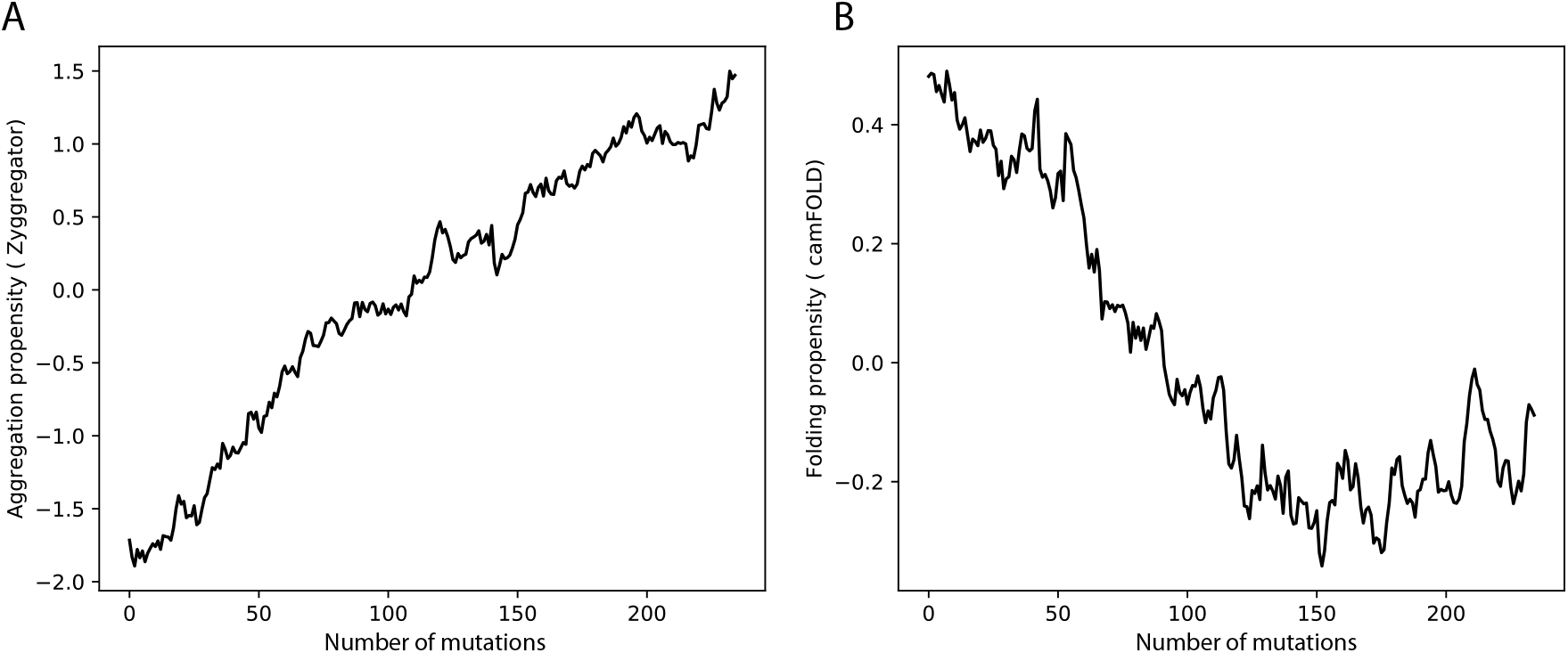
Aggregation and folding propensities changes upon amino acid mutations in human proteome. A) The aggregation propensity increases while the protein accumulate random mutations. By contrast, B) the folding propensity decreases when the protein is subjected to mutation.

### C. Aggregation as a context-dependent constraint on protein evolution

To address the question on how much the environment influences the role that aggregation has in the evolution of proteins, we analyzed data from two different molecular evolution experiments. In the first experiment, TDP-43 was heterologously expressed in *S. cerevisiae* and the toxicity of variants collected. In the second experiment, variants of TEM-1 beta-lactamase were selected in *E. coli* while antibiotics were given to the cell culture.

In this analysis we used the ROC analysis to measure the strength of the link between the toxicity and the aggregation propensity [2]. The ROC is highly appropriate as it allows to simultaneously analyse the true and false positive rates.

#### Cell fitness and aggregation propensity of TDP-43 mutants

In the case of TDP-43 expression in *S. cerevisiae*, we divided the experimental sets in two partitions of equal size corresponding to the highest (positive) and lowest (negative) fitness values associated with the mutants. As shown in Figure 3, the Area Under the ROC Curve (AUC) rises progressively with the high vs low fitness mutations score. Our result indicates that the aggregation is an excellent predictor of fitness, especially for those mutants that have the strongest experimental signal (AUC of 0.78 or more). In the insets of Figure 3 we show statistics for the aggregation propensities of single and double mutations associated with the highest and lowest fitness scores (corresponding to AUCs of 0.78 and 0.77, respectively).

**FIG. 3:**
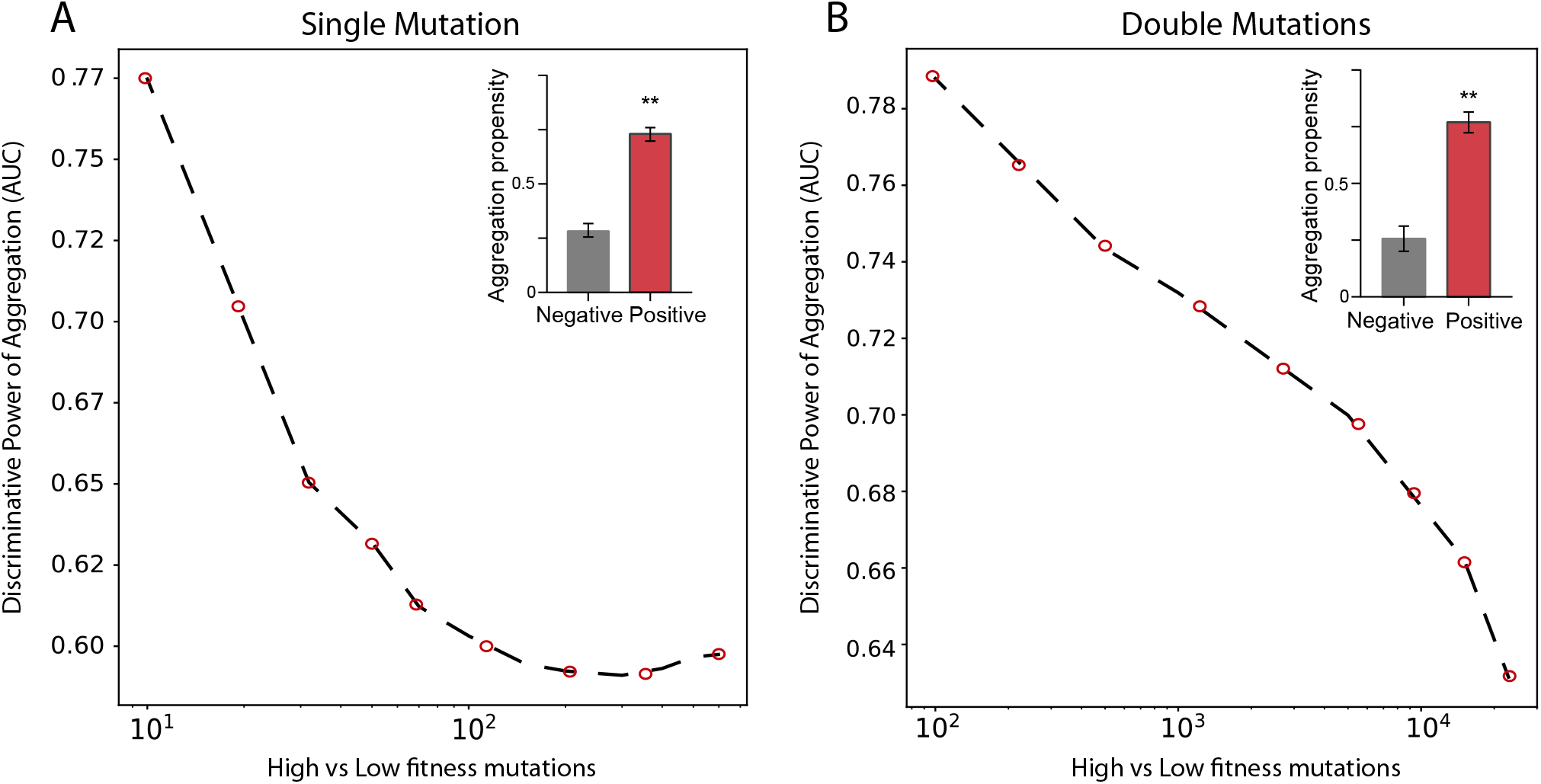
Deep mutational scanning experiments of TDP-43 mutants expressed in *S. cervisiae*. AUC scores computed on the top and bottom sub-data sets. Panel A and B show the rise of the AUC values as function of the high vs low fitness data sets for single and double mutations, respectively.In the insets, we show the aggregation propensities of mutations associated with the highest (top 100) and lowest (bottom 100) fitness mutations (**, p-value*<*0.01, Kolmogorov-Smirnoff test)

We then swapped the negative set with a list of mutants generated randomly (Figure 4; Materials and Methods). We observed that the AUC increases monotonically with the signal strength encoded in the aggregation propensity of the high fitness score mutants (statistics of the mutations associated with the 100 highest vs 100 random mutations are shown in Figure 4B). This result indicates that the negative and random sets are interchangeable for our analyses.

**FIG. 4:**
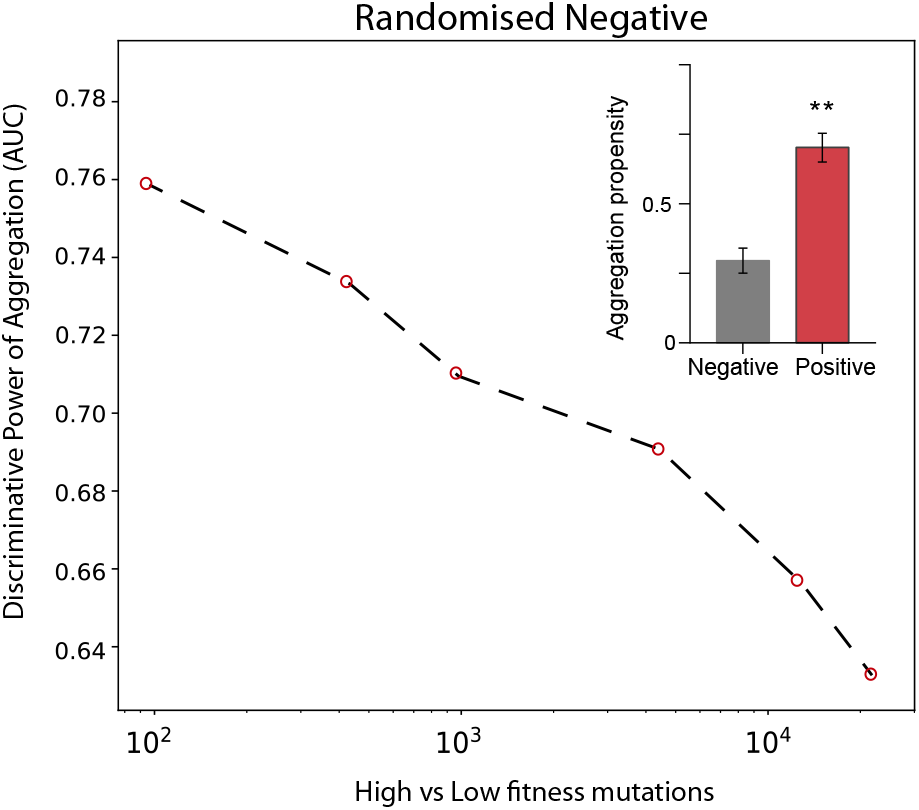
Deep mutational scanning experiments of TDP-43 mutants expressed in *S. cervisiae* using a random negative set. The AUC values increase progressively with the experimental signal strength. Double mutants are analysed. In the insets, we show the aggregation propensities of mutations associated with the highest (top 100) and random (100 cases) fitness scores (**, P-value*<*0.01, Kolmogorov-Smirnoff test).

### Cell fitness and aggregation propensity of TEM-1 beta-lactamase mutants

In the case of TEM-1 beta-lactamase in *E. coli*, only the positive fitness set was available from the experiments. To generate a negative set following the approach used for TDP-43, we first analysed the number of mutations and sequence lengths of variants. We observed that the number of mutations increases on average with the generations (Figure 5, panels A and B). Focusing on the variants that have the same length as the reference TEM-1 beta-lactamase, we generated the negative set respecting the frequencies of mutations occurring at each cycle. In Figure 5C we show the values corresponding to the AUC computed using the aggregation propensities for the high (positive) vs low (negative) fitness score mutants. There is a clear trend towards a decrease of the AUC score (i.e., increase of 1-AUC) with the number of mutations and evolutionary cycles (generation). Thus, differently from TDP-43, we observed that the aggregation propensity decreases. In Figure 5D we show that the aggregation propensity of mutants associated with high fitness score decreases progressively through evolutionary cycles. In this analysis, we selected variants by ranking the number of mutations occurring at each evolutionary cycle and compared their aggregation propensities with their random counterpart (100 sequences for each set; the AUCs are 0.6, 0.65 and 0.70, respectively).

**FIG. 5:**
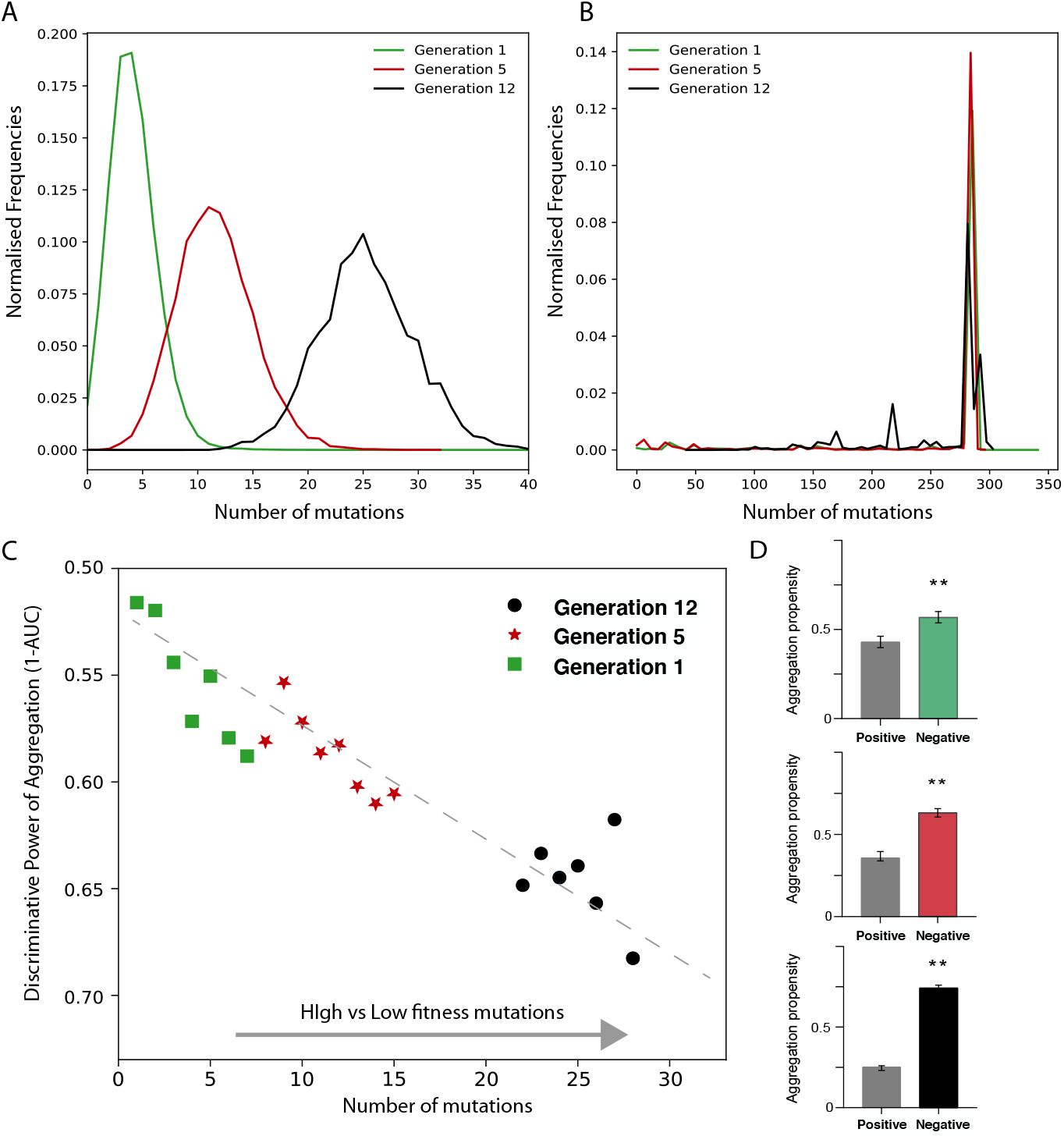
Deep scanning experiments of TEM-1 beta-lactamase mutants expressed in *E. coli*. Distribution of A) mutation number and B) Distribution of sequence length for different generation of evolution. C) AUC value as a function of high vs low fitness data sets (data computed only for sub data set with at least 1000 data points). D) Aggregation propensities of mutations associated with the highest (top 100) and random (100 cases) fitness scores for generation 1,5, 12 (**, p-value*<*0.01, Kolmogorov-Smirnoff test). The colors match the points in the panel D.

## IV. DISCUSSION

This study aimed to investigate the influence of protein aggregation on molecular evolution. Mutational scanning approaches have been previously used in the literature and demonstrated to provide a powerful tool to investigate the ability of a protein to aggregate and cause toxicity [27]. We also have previously observed that, under physiological conditions, expression levels are fine-tuned to the specific protein solubilities [8]. This finding suggests that mutations altering the aggregation propensity can impact cell fitness and thus influence evolution [7, 8].

The computational analysis presented in this paper revealed that aggregation and folding propensities anti-correlate at the proteomic level. These results agree with and expand studies that had suggested similar conclusions from completely different premises [22, 28]. Our data indicate that, for the large majority of cases, naturally-occurring protein sequences have been optimized by evolution to have lower aggregation and higher folding propensities [14]. Our calculations predict that random mutations render proteins more prone to aggregate at the expenses of their folding ability, even though this variability could help exploring new functionalities depending on the cellular context.

To estimate the impact of aggregation in different cellular contexts, we analysed molecular evolution experiments in which human TDP-43 and TEM-1 beta-lactamase mutants were expressed in *S. cerevisiae* [23] and in *E. coli* [24] respectively. Notably, TDP-43 contains *>* 30% of structural disorder in the C-terminus [3], whereas TEM-1 beta-lactamase has a well-defined native state [24]. When prion-like regions, as in the case of TDP-43 [23], are not protected by the native fold, aggregation takes place because the exposure to the solvent favours self-assembly [2, 3, 29]. For TDP-43 mutants, both negative and positive fitness scores were directly compared with the aggregation propensities. By contrast, for TEM-1 beta-lactamase, we had only positive fitness variants selected through the evolutionary process, which made the comparison with a negative set not directly possible. To carry out the analysis, we artificially generated a negative set as close as possible to the experimental one.

Our analysis revealed two quite different behaviors. With TEM-1 beta-lactamase, the system was under antibiotic selection pressure. Thus, the protein had to remain active to protect against penicillins and avoid aggregation. The variants were in fact selected by evolution to maintain enzyme activity and thus antibiotics resistance while minimizing aggregation [24]. While the negative set maintains a relative high aggregation propensity through the evolutionary cycles, the positive set significantly reduces it, showing existence of a purifying pressure. Interestingly, it has been previously observed that M180T, E195D, L196I, and S281T were the most frequent mutations in the system. These residues are all exposed and their mutations should, with the exception of the well documented M180, only have minor effects on protein fold as indicated by visual inspection and semi-quantitative analysis with the DUET software, a web server for an integrated computational approach to study missense mutations in proteins [30]. Thus, these mutations would represent a “neutral” passage that do not impinge on fitness.

With TDP-43, we observed the opposite trend. We have recently stressed the importance of protein interactions with chaperones to prevent aggregation [10, 13, 15]. Expression of a protein in a heterologous system will lead by definition to a situation in which natural interactions cannot occur [3]. Accordingly, we observed that TDP-43 has a high toxicity in yeast, as expected for a protein that has a strong tendency to aggregate also in its natural host [23]. In this case, we observed that mutations that increased hydrophobicity and aggregation also decreased toxicity by eliminating from solution proteins that could not have functional partners. This hypothesis would explain why solid aggregates formed by TDP-43 A238V, Q360Y and G335I mutants are non-toxic, while the soluble W334K, M332K and A328P have a higher toxicity, as described in Bolognesi ed al. [23]. This would also be consistent with the generally accepted idea that soluble aggregates are more toxic than the insoluble end-points [31], although in this work we did not analyse liquid-liquid phase separations.

It is worth mentioning that the TDP-43 mutational library was designed to target a specific region of the protein, the intrinsically unstructured C-terminus. This meant no evolution pressure to preserve structure. Accordingly, our results showed that the aggregation propensity of TDP-43 increased with the fitness score.

The two scenarios indicate a clear ambivalence that protein aggregation can acquire depending on the system in which it occurs and suggest aggregation as an important constraint for evolution. Our approach demonstrates how different selective pressures may completely change the effect of aggregation on a specific system: an exogenous protein with no functional role, such as TDP-43 in *S. cerevisiae* is less toxic if compartimentalised in an aggregate, while an endogenous proteins, such as TEM-1 beta lactamase in *E. coli*, is functional when aggregation is avoided. Further analyses will be important to shed light onto the selective pressure against aggregation in specific cellular environments. Indeed, it has been previously suggested that the volume of the organelles imposes stringent limitations on protein evolution to avoid high aggregation propensities in confined compartments [32, 33]. This has implications for protein variability in different organisms [34]. In line with this observation, it should be mentioned that poly-anionic molecules such as RNA can reduce aggregation propensity [35] and [36]. On the contrary, mutations that abolish RNA-binding in these proteins can force them to change their cellular localization, accumulate and aggregate. This is the case of FUS and TDP-43 [35, 37]. Making large-scale predictions of protein aggregation can reveal important properties of biological systems, as demonstrated by the elegant study by Khodaparast et al. [38] who showed that identifying insoluble regions of a protein can be instrumental to block the process of bacterial infection.

In the future, it will be interesting to investigate the role of specific molecular partners, such as RNA, in regulating aggregation. Computational approaches such as the one presented in this study [39, 40] will be key to achieve a more complete understanding of molecular evolution, and to increase our ability to manipulate proteins for specific purposes.

## V. ACKNOWLEDGEMENTS

The authors would like to thank Dr. Mattia Miotto, Dr Lorenzo Di Rienzo, Dr Tommaso Muto and Dr. Claudia Giambartolomei for discussions. The research leading to these results was supported by the Dementia Research Institute (REI 3556) and AlzUK (ARUK-PG2019B-020), European Research Council (RIBOMYLOME 309545 and ASTRA 855923), the H2020 projects IASIS 727658 and INFORE 25080, the Spanish Ministry of Economy and Competitiveness BFU2017-86970-P as well as the collaboration with Peter St. George-Hyslop financed by the Wellcome Trust.

