## Supplementary Materials for "Aggregation is a context-dependent constraint on protein evolution" (from another version of this preprint: version 1, 2021-05-11)

### I. HUMAN PROTEOME

In Figure 1 we report the distribution of the sequences length and the frequencies of amino acids for the human proteome. The histograms in Figure 2 show four curves, two for the aggregation and two for the folding propensity, computed for uniformly and weighted distributed amino acids (distribution shown in Figure 1 right panel) . We found that the folding propensity is reduced through the process of randomization while the aggregation increase on average. This is consistent in both randomizations used.

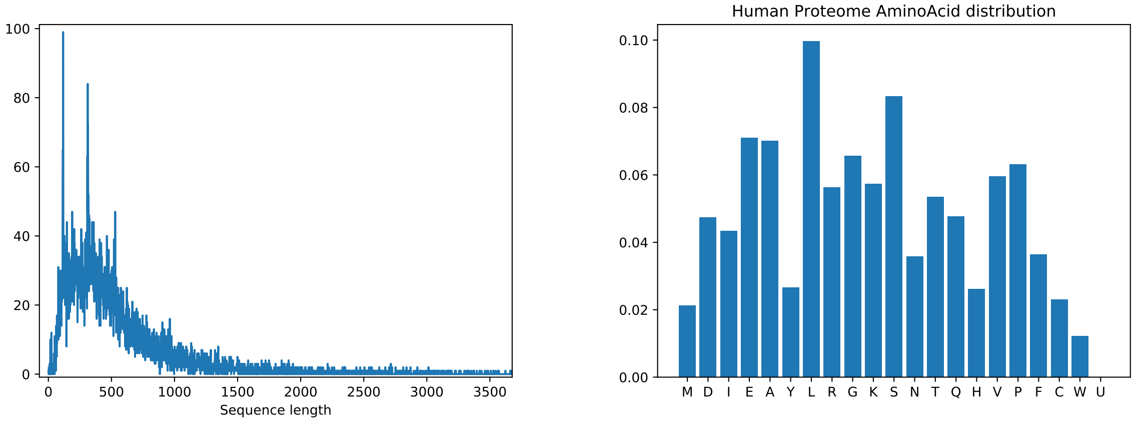


FIG. 1: Statistical analysis of the human proteome. On the left side we report the distribution of the lengths of protein sequences, and on the right the amino acid frequencies.

### II. TDP-43 EXPRESSION IN S. CEREVISIAE

Analysis carried out with the Clever Machine approach reveals that aggregation strongly discriminates high vs low fitness mutants (5000+5000 instances analysed):

`http://crg-webservice.s3.amazonaws.com/submissions/2020-09/295992/output/index.html?unlock=`

e24ba9a2ff.

For different high vs low fitness scores, we report the performances of the linearized Zyggregator algorithm computed using the Area under the Receiver Operating Characteristics Curve (AUC of ROC; Figure 3). In Figure 4 we show the predicted aggregation propensity for the experimental negative set and its random version (single and double mutations).

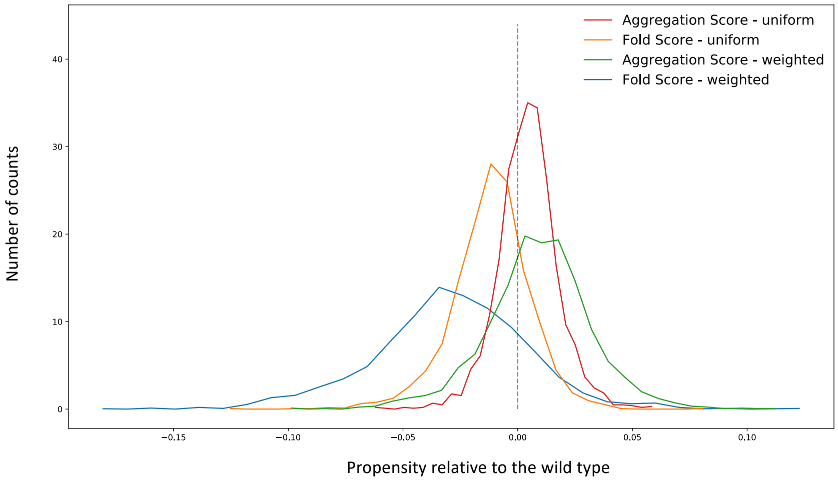


FIG. 2: Difference between aggregation and folding propensity between the mutated protein and the WT one. Each final protein has 500 random mutations respect to the WT one. The histograms is computed over 10 000 random protein of the human proteome. The random is computed in two ways: considering the amino acid distribution uniform or using the human proteome amino-acid frequency. The results do not change .

### III. TEM BETA-LACTAMASE EXPRESSION IN E. COLI

AUC for all the sub datasets regarding the TEM Beta-Lactamase experiment. In Figure 5 we show how we filtered the data taking into account only the AUC that have been computed for significantly populated datasets. Datasets of a small size and with an higher number of mutations respect to the mean of the generation class are discarded. Sequences with higher number of mutations are more prone to have a lower fitness propensity, as shown in Figure 5.

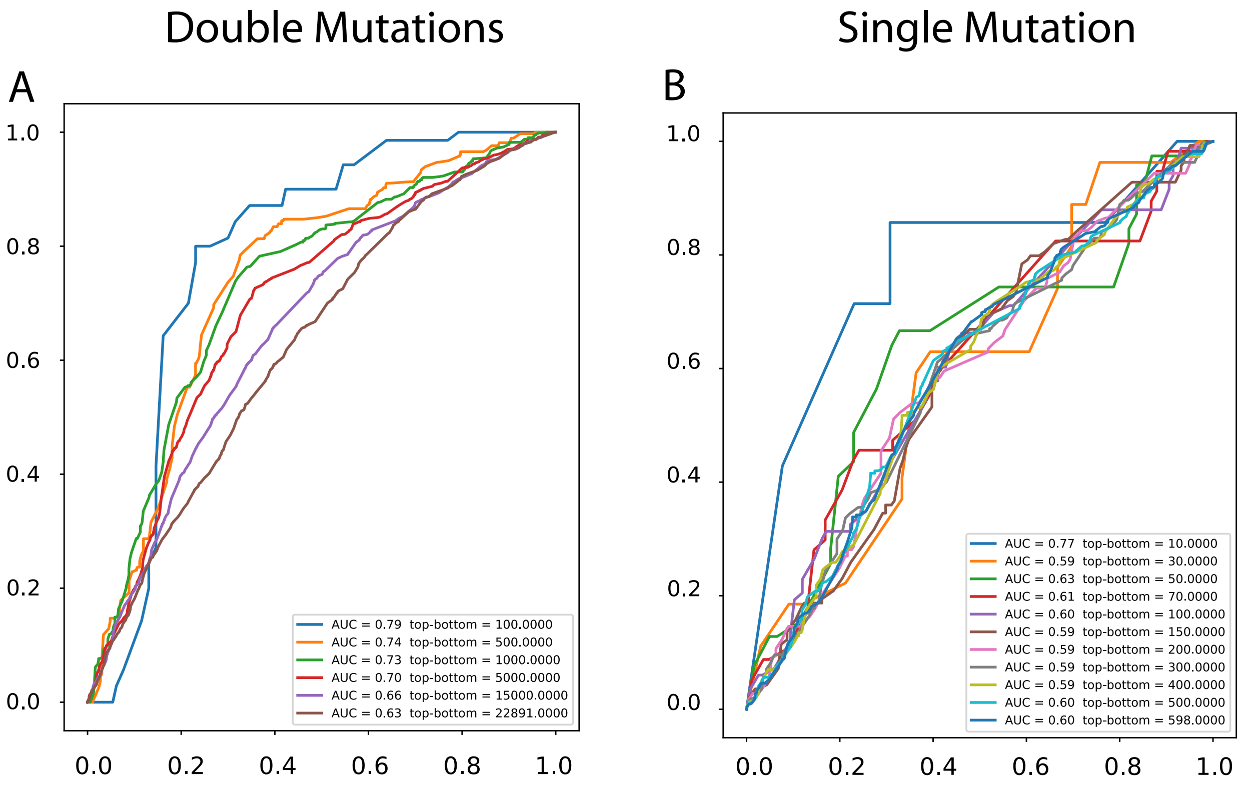


FIG. 3: Aggregation propensity of TDP-43 mutants discriminate high vs low fitness scores. The ROC are curves computed for different high vs low fitness mutations. Panel A shows double mutations dataset and B single mutations mutation.

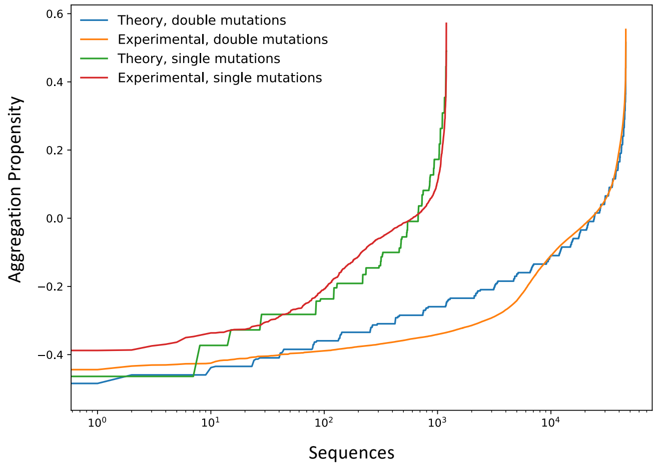


FIG. 4: Aggregation propensities of experimental and theoretical mutants and related aggregation propensities.

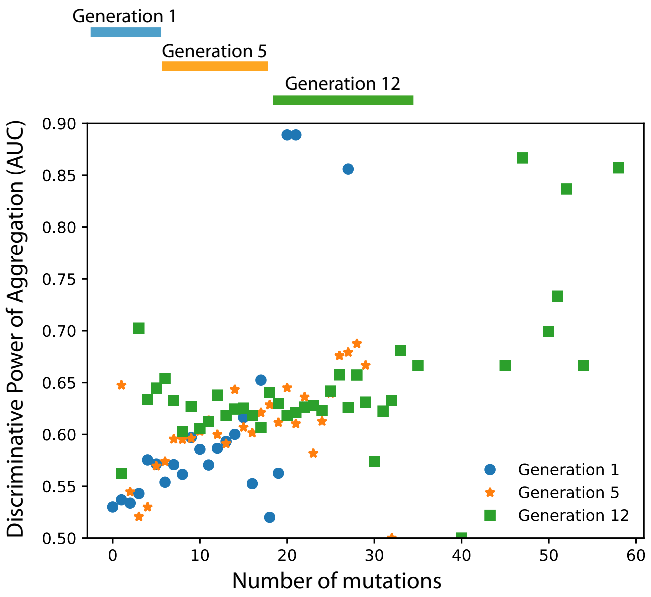


FIG. 5: AUC computed for TEM Beta-Lactamase mutants. In the analysis we selected only points belonging to datasets of significant size (mutational range highlighted with colored bars; Materials and Methods).
